# Sympathetic Signals Promote Immunosuppressive Neutrophils in Lung Tumors

**DOI:** 10.1101/2024.02.15.580434

**Authors:** Jiaqi Liu, Mantang Qiu, Wenxiang Wang, Ying Cao, Fan Yang, Kezhong Chen, Jian Chen, Jing Yang

**Author notes:** Corresponding authors (K.C.), (J.C.), (J.Y.). Authors contributed equally (Jiaqi Liu, Mantang Qiu).

## Abstract

It has become recognized that the nervous system can significantly influence cancer prognosis. However, the pathophysiological mechanism underlying neural modulation of the tumor immune microenvironment remains incompletely understood. Here, we exploit the advanced 3D imaging technique to reveal the frequent presence of sympathetic inputs in human non-small-cell lung cancer. Also, this spatial engagement of sympathetic innervations similarly occurs in the mouse models of lung tumors. Of importance, the local ablation of sympathetic signals strongly suppresses cancer progression. We then identify a neutrophil subtype uniquely present within lung tumors, whose immunosuppressive features are directly promoted by the sympathetic neurotransmitter norepinephrine via the β2-adrenergic receptor (Adrb2). In addition, norepinephrine stimulates the specific type of tumor cells to recruit such immunosuppressive neutrophils. Accordingly, we show that the Adrb2 antagonist effectively potentiates the anti-PD-L1 immunotherapy. Together, these results have elucidated a critical aspect of sympathetic signals in designating the immune microenvironment of lung tumors.

## Introduction

The immune microenvironment has a determinant role in tumor onset, growth, and metastasis, and it has been broadly realized that cancer cells may evolve diverse mechanisms to suppress or evade the body’s antitumor immunity ^1–5^. Among them, tumor cells can express immune checkpoint ligands, e.g., CD274 (commonly known as PD-L1), to block the function of cytotoxic CD8^+^ T cells ^6–8^. In addition, tumor-associated macrophages ^9–11^ or regulatory T cells ^12,13^ are often recruited to produce specific immunosuppressive signals, e.g., arginase 1, interleukin 10, and galectins. Such tumor-infiltrating immune cells also elicit angiogenetic cues, e.g., vascular endothelial growth factors, to facilitate cancer proliferation and metastasis. Therefore, a comprehensive understanding of the tumor immune microenvironment holds the promise of effective treatments and even a cure for dreadful human cancers ^14–17^.

Accumulating evidence has suggested that neural signals significantly influence the prognosis of various cancer types. Indeed, the emerging frontier of cancer neuroscience has increasingly garnered research attention in the past years ^18–20^. For instance, the presence of sympathetic or parasympathetic innervations in prostate tumors correlated with poorer survival of patients ^21^. Conversely, the incidence of prostate cancer decreased in the male population inflicted by spinal cord injuries that would conceivably abrogate the efferent sympathetic or parasympathetic actions ^22^. Similarly, in the women taking β-adrenergic receptor-antagonizing drugs (i.e., beta blockers) that chronically attenuated sympathetic signals, the rates of metastasis or mortality became significantly lower for breast cancer ^23,24^ or ovary cancer ^25^. Notably, such clinical observations on the involvement of the body’s nervous system in cancers can be recapitulated in mouse models, which help provide more mechanistic insights. For example, the local ablation of sympathetic or parasympathetic innervations in the mouse prostate revealed their disparate roles in tumorigenesis or tumor progression ^21^. Also, the surgical or pharmacological blockage of neural innervations in the mouse stomach suppressed the Wnt-mediated cell proliferation, thus ameliorating the onset and growth of gastric cancer^26^. In addition, the beta-blocker treatment in mice slowed the metastasis of breast cancer by decreasing the infiltration of tumor-associated macrophages ^27^. Further, inhibiting the neurotransmitter glutamate signal via the N-methyl-D-aspartate receptor (NMDAR) reduced the invasiveness of the mouse pancreatic neuroendocrine tumor or pancreatic ductal adenocarcinoma ^28^. Despite those research advances, the nervous system’s complex crosstalk with cancers calls for more in-depth investigations.

Clinical analyses reported that the prognosis of non-small-cell lung cancer (NSCLC) tended to deteriorate in people with symptoms of anxiety or depression ^29–32^, implicating neural signals in modulating the disease course. Also, the use of beta-blockers improved the survival outcomes of NSCLC patients receiving radiotherapy ^33^, chemotherapy ^34^, or immunotherapy ^35^, though such beneficial effects are still being debated ^36,37^. However, the pathophysiological mechanism underlying this potential sympathetic regulation of lung cancer is incompletely understood. In particular, recent studies revealed the heterogeneous neutrophil subtypes in human NSCLC, suggesting their critical role in establishing the immune microenvironment ^38,39^. Whether sympathetic signals may instruct the specific function of neutrophils within lung tumors is entirely uncharted.

## Results

As an essential entry point of this study, we sought to assess whether there would be a direct link between lung tumors and the nervous system. Given clinical observations on the potential beneficial roles of beta blockers in human NSCLC, we focused on the involvement of sympathetic inputs. Notably, conventional immunohistochemistry methods based on tissue sectioning have intrinsic limitations in assessing 3D neural distributions, and such technical shortcomings may lead to incomplete or even controversial results. To solve this conundrum in the field, colleagues and we have developed a series of three-dimensional (3D) imaging techniques in the past decade, enabling the comprehensive assessment of neural distributions in various intact mouse or human organs, including the lung ^40–43^. Therefore, we pursued this advanced imaging approach to examine sympathetic inputs in human NSCLC samples. The unsectioned tumors were processed by the established procedure of tissue permeabilization and whole-tissue immunolabeling of tyrosine hydroxylase (TH), a specific marker for sympathetic axons. The immunolabeled NSCLC tumors were then subjected to optical clearing, which rendered the samples almost transparent for lightsheet microscopy (Fig. 1A). Of importance, we observed abundant TH-positive sympathetic axons within the human NSCLC tumors by the 3D reconstruction of lightsheet imaging (Fig. 1B). In addition, the optical sections of lightsheet microscopy unequivocally demonstrated that those sympathetic innervations were projecting inside of those solid tumors (Fig. 1C). Indeed, TH-positive sympathetic axons were observed in 18 out of 20 patient samples (Table S1), suggesting it as a common feature of human NSCLC.

**Figure 1.**
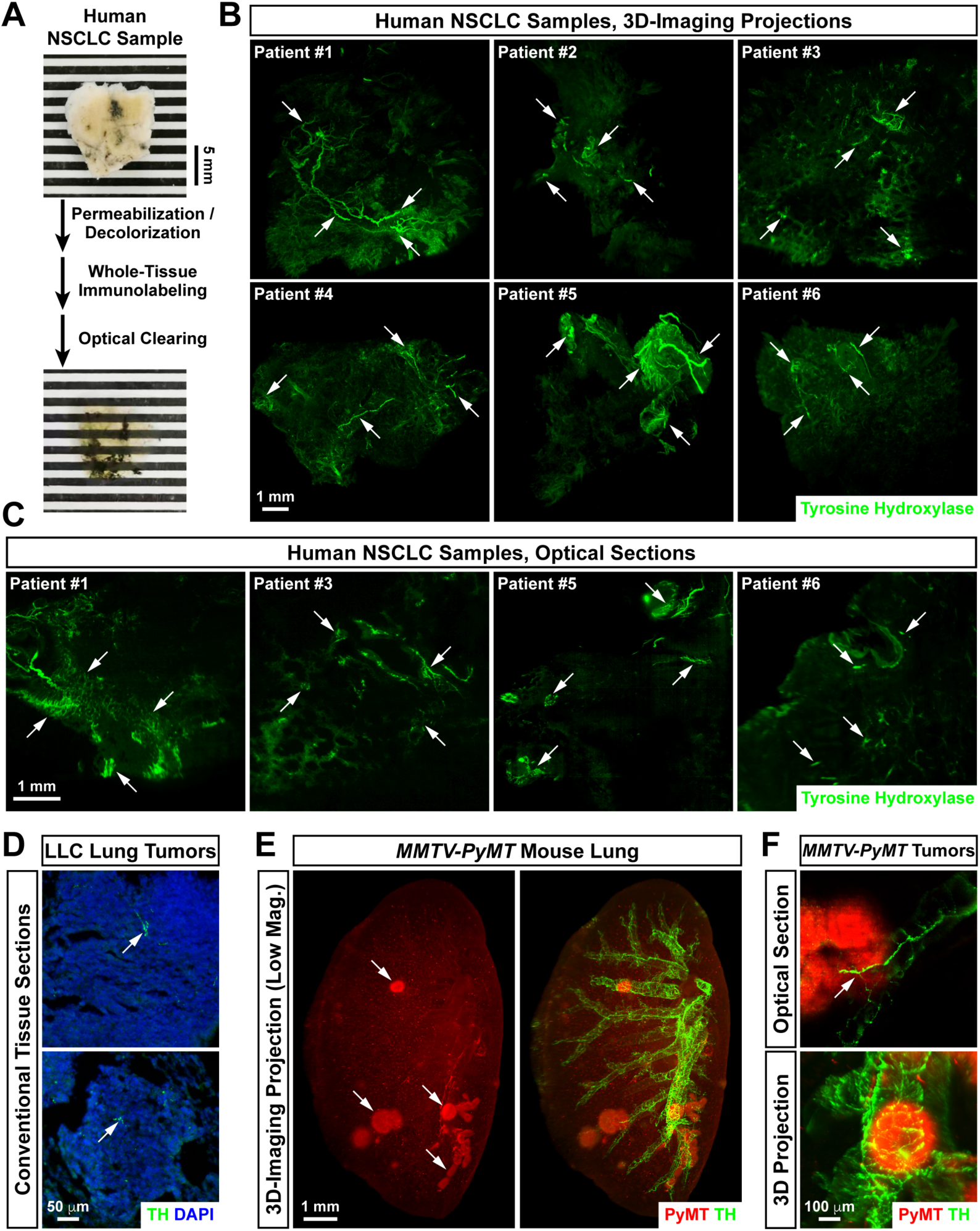
Spatial engagement of sympathetic innervations with human or mouse lung tumors. **(A to C)** Presence of sympathetic inputs in human NSCLC. The unsectioned human NSCLC samples (patient information summarized in Table S1) were processed through tissue permeabilization and decolorization, whole-tissue immunolabeling of tyrosine hydroxylase (TH), and optical clearing for lightsheet 3D imaging. **(A)** Representative images of a human NSCLC sample before and after the processing steps. **(B and C)** Representative 3D-projection images at 1.26× magnification **(B)** or optical sections at 2.0× magnification **(C)** of the lightsheet imaging were shown. Arrows exemplify the TH-positive sympathetic axons within human NSCLC samples. **(D)** Sympathetic innervations in the lung tumors of LLC mouse model. Representative images of the lung tumors at 30-day post-inoculation processed by conventional tissue sectioning and TH-immunostaining were shown. Arrows denote the TH-positive sympathetic axons in LLC lung tumors. **(E and F)** Sympathetic innervations in the metastatic lung tumors of *MMTV-PyMT* mouse model. The intact, unsectioned lung tissues (left lobe) of *MMTV-PyMT ^+/-^*female mice at 7 weeks after the onset of primary breast cancer were processed through the whole-tissue co-immunolabeling of polyomavirus middle T antigen (PyMT, red) and TH (green). **(E)** Representative 3D-projection images at 1.26× magnification of the lightsheet imaging were shown. Arrows exemplify the PyMT-positive metastatic lung tumors. **(F)** Representative 3D-projection image (upper panel) and optical section (lower panel) of metastatic lung tumors at 8.0× magnification of the lightsheet imaging were shown. The arrow denotes the TH-positive sympathetic axons within a lung tumor.

We looked into whether sympathetic innervations would be similarly detected in mouse lung tumors. First, we examined the standard model of orthotopic lung tumors of Lewis lung carcinoma (LLC) cells. For comparison with lightsheet 3D imaging, LLC lung tumors were subjected to conventional tissue sectioning. TH-immunostaining exhibited the sympathetic axons inside tumors, though they appeared more fragmented on tissue sections (Fig. 1D). Next, we investigated whether sympathetic innervations might also be present in metastatic lung tumors. To this end, we utilized the *MMTV-PyMT* mouse model in which the onset of primary breast cancer would be followed by lung metastases. The intact, unsectioned lung tissues of tumor-bearing *MMTV-PyMT ^+/-^* mice were subjected to the whole-tissue co-immunolabeling of TH together with the oncogenic PyMT protein expressed in cancer cells. Strikingly, a spatial engagement of TH-positive sympathetic axons with PyMT-positive metastatic lung tumors was revealed by 3D imaging (Fig. 1E). High-magnification 3D reconstruction or the optical section of lightsheet microscopy confirmed that the sympathetic axons were localized insides or ensheathing tumors (Fig. 1F). These results have demonstrated that reminiscent to those observed above in human NSCLC samples, sympathetic inputs are connected to mouse lung tumors.

We set out to explore the functional relevance of sympathetic signals in lung tumors. The mice were intranasally treated with 6-hydroxydopamine (6-OHDA) to ablate local sympathetic innervations in the lung, which was verified by the whole-tissue TH-immunolabeling and 3D imaging (Fig. 2A and 2B). The key advantage of this pharmacologic approach is that it would not affect sympathetic structures in other organs, e.g., the thymus, cervical lymph nodes, spleen, or femoral bone marrow, as we previously reported ^43^. Also, the loss of sympathetic signals did not affect immune cell populations relevant to tumor immunity in the lung, e.g., CD8^+^ T cells or neutrophils (Fig. 2C, see below). In addition, there were no detectable changes in the plasma levels of oxygen or glucose of 6-OHDA-treated mice (Fig. S1A), ruling out an overall impact on metabolism. However, this specific ablation of sympathetic signals strongly suppressed tumor growth in the LLC lung cancer model (Fig. 2D and 2E). Meanwhile, an increase of CD8^+^ T cells occurred in the lung tumors of 6-OHDA-treated mice (Fig. 2F and 2G), suggesting an enhancement of antitumor immunity. In parallel, we tested the effect of sympathetic ablation in the *MMTV-PyMT* cancer model. As expected, the specific destruction of sympathetic distribution in the lung did not affect the progression of primary breast tumors (Fig. S1B). In contrast, 3D imaging clearly showed an inhibition of the onset and growth of PyMT-positive metastatic lung tumors (Fig. 2H and 2I). Similar to that in the LLC cancer model, sympathetic ablation boosted CD8^+^ T cells in *MMTV-PyMT* lung tumors (Fig. 2J and 2K). These results have elucidated that local sympathetic inputs can exert a stimulative effect on lung tumors.

**Figure 2.**
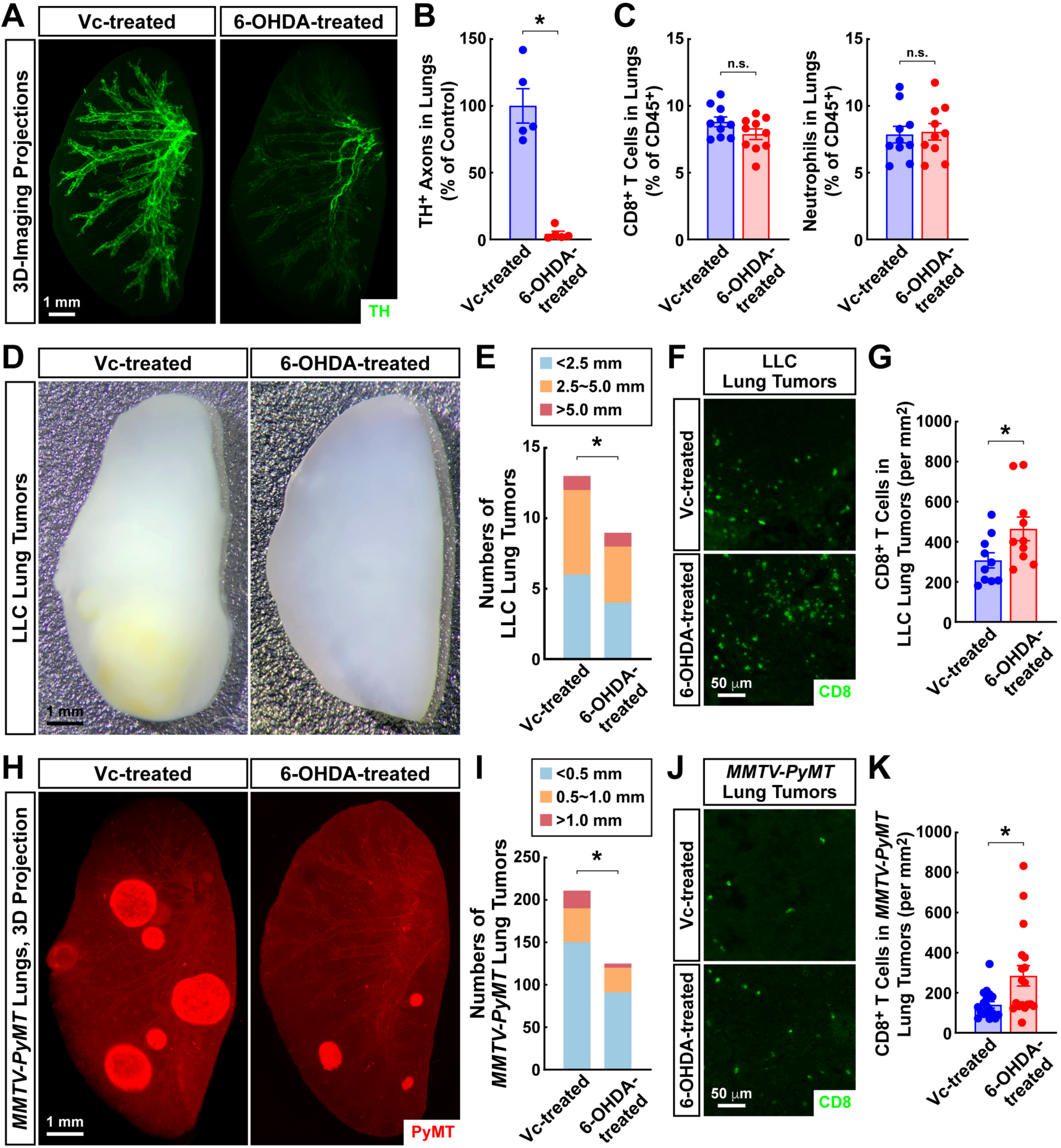
Ablation of local sympathetic signals in the lung inhibits tumor growth. **(A to C)** Pharmacologic ablation of sympathetic innervations in the lung. The wild-type mice were subjected to the intranasal treatment of 6-OHDA or control vehicle [PBS containing 0.01% vitamin C (Vc)]. **(A and B)** The intact, unsectioned lung tissues (left lobes) of Vc-treated or 6-OHDA-treated mice were processed for the whole-tissue TH-immunolabeling. **(A)** Representative 3D-projection images at 1.26× magnification of the lightsheet imaging were shown. **(B)** TH-positive sympathetic axons in the lung tissues were quantified. n = 5, mean ± SEM, * *p* < 0.05 (Student’s *t*-test). **(C)** CD8^+^ T cells and neutrophils in the lung tissues of Vc-treated or 6-OHDA-treated mice were examined by the FACS analyses. n = 10, mean ± SEM, n.s., not significant (Student’s *t*-test). **(D to G)** Sympathetic ablation inhibits tumor growth in the LLC cancer model. The wild-type mice were intranasally treated with 6-OHDA or control vehicle and then inoculated with LLC cancer cells. **(D)** Representative images of the lung tumors of Vc-treated or 6-OHDA-treated mice at 30 days post-inoculation were shown. **(E)** Numbers and sizes of the lung tumors of Vc-treated or 6-OHDA-treated mice were measured. 9 mice per condition, * *p* < 0.05 (two-way ANOVA test). **(F)** Representative images of the lung tumors processed by conventional tissue sectioning and CD8-immunostaining were shown. **(G)** Numbers of CD8^+^ T cells in the lung tumors of Vc-treated or 6-OHDA-treated mice were quantified. n = 10, mean ± SEM, * *p* < 0.05 (Student’s *t*-test). **(H to K)** Loss of sympathetic signals suppresses metastatic lung tumors in *MMTV-PyMT* mouse model. *MMTV-PyMT ^+/-^* female mice at 1 week after the onset of primary breast cancer were intranasally treated with 6-OHDA or control vehicle. **(H and I)** The intact lung tissues (left lobes) of Vc-treated or 6-OHDA-treated *MMTV-PyMT ^+/-^*mice at 9 weeks after the onset of primary breast cancer were processed through the whole-tissue TH-immunolabeling. **(H)** Representative 3D-projection images at 1.26× magnification of the lightsheet imaging were shown. **(I)** Numbers and sizes of the metastatic lung tumors of Vc-treated or 6-OHDA-treated *MMTV-PyMT ^+/-^*mice were counted. 32 mice per condition, * *p* < 0.05 (two-way ANOVA test). **(J)** Representative images of the metastatic lung tumors processed by conventional tissue sectioning and CD8-immunostaining were shown. **(K)** Numbers of CD8^+^ T cells in the metastatic lung tumors of Vc-treated or 6-OHDA-treated *MMTV-PyMT ^+/-^*mice were quantified. n = 18, mean ± SEM, * *p* < 0.05 (Student’s *t*-test).

We went on to determine how sympathetic signals in the lung influence tumor progression. Studies by our colleagues and us have begun to document the diverse roles of neutrophils in shaping the tumor immune microenvironment ^38,39,44,45^. Indeed, our in-depth analyses of the published scRNA-seq dataset have identified the presence of immunosuppressive neutrophils in the LLC lung tumors [Fig. S12 in ^45^]. As a unique feature of those neutrophils, they were only present inside tumors but not in peripheral lymphoid organs such as the spleen of tumor-bearing mice. Importantly, this distinct neutrophil subtype expressed several immunosuppressive genes such as *Cd274* (commonly known as PD-L1). Also, those immunosuppressive neutrophils could produce myeloid recruitment-related genes, including *Ccl3*, *Ccl4*, and *Cxcl2*, to designate the immune microenvironment of LLC lung tumors.

To examine whether immunosuppressive neutrophils would be present in metastatic lung tumors, we first performed the scRNA-seq analyses of total immune cells in the lungs of *MMTV-PyMT ^+/-^* mice. Compared to *MMTV-PyMT ^-/-^* wild-type littermates, there was a profound accumulation of neutrophils among the immune cell populations defined in the scRNA-seq dataset (Fig. 3A). This neutrophil response could be verified by the FACS analyses, showing approximately 4-fold increase of neutrophils in the lungs of *MMTV-PyMT ^+/-^* mice (Fig. 3B). We next pursued the scRNA-seq analyses focusing on neutrophils in the peripheral blood, paracancerous lung tissues, and metastatic lung tumors of *MMTV-PyMT ^+/-^*mice. Significantly, a distinct neutrophil subtype was exclusively detected in metastatic lung tumors but not the blood or paracancerous lung tissues of tumor-bearing mice (Fig. 3C and 3D). Those neutrophils likely originated from the differentiation of neutrophil populations encountered in the blood circulation or normal lung tissues (Fig. 3E). Similar to the counterpart in the LLC cancer mode, this unique neutrophil subtype in *MMTV-PyMT* lung tumors simultaneously expressed multiple immunosuppressive genes such as *Cd274* and *Cd81*, together with several myeloid recruitment-related factors (Fig. 3F and Fig. S2). Such a signature expression of *Cd274* by immunosuppressive neutrophils in LLC or *MMTV-PyMT* lung tumors was validated by the FACS analyses (Fig. 3G, Fig. S3A, and Fig. S3B). As a distinct morphological feature of neutrophils, their nuclei are heavily segmented into multiple lobes. We noted that despite the differential subtypes, neutrophils within LLC or *MMTV-PyMT* lung tumors maintained the appearance of nuclear segmentation (Fig. S3C and S3D), serving as a verification of their cellular identity.

**Figure 3.**
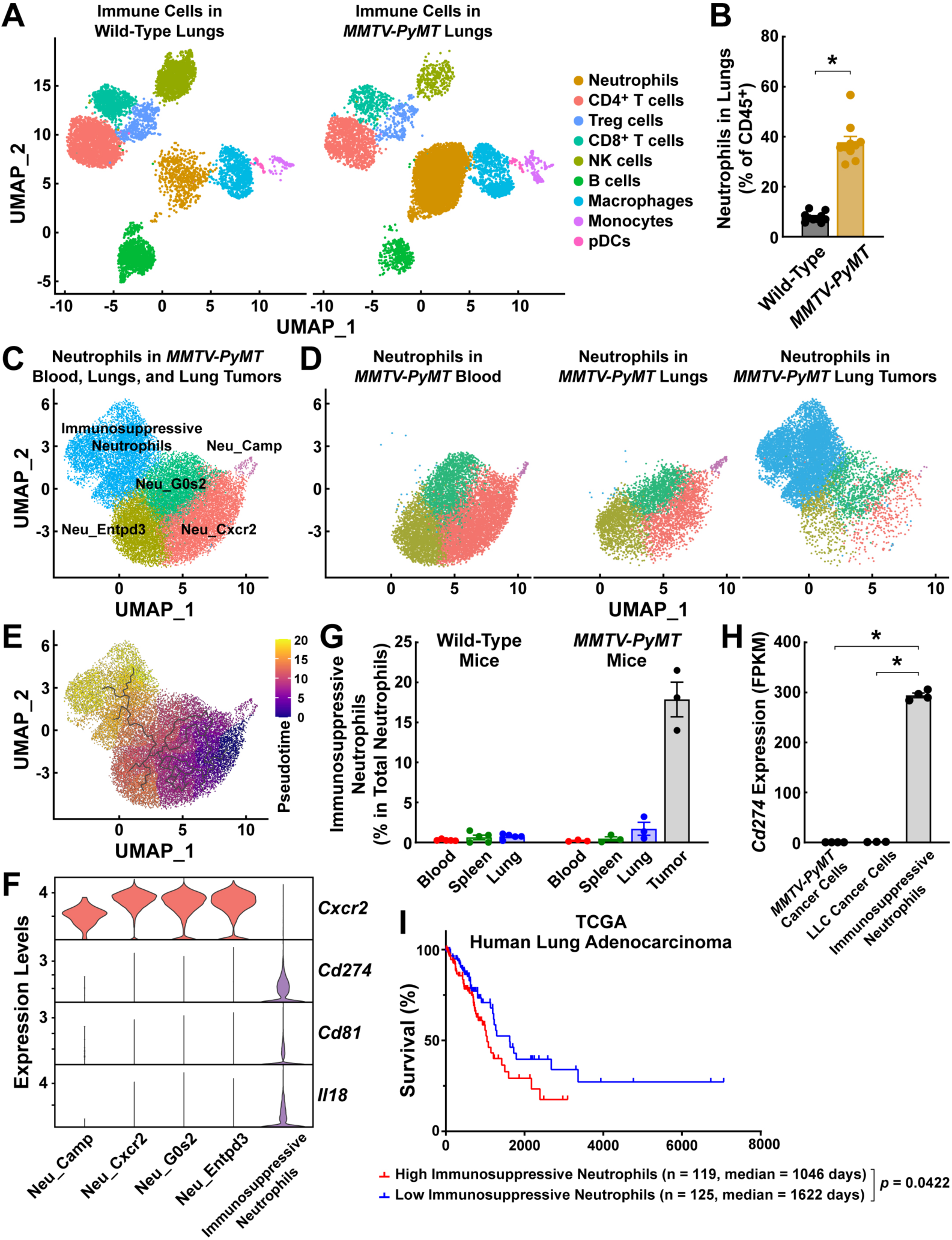
Immunosuppressive neutrophils are uniquely present within lung tumors. **(A and B)** Neutrophil responses in the lung tissues of *MMTV-PyMT* mouse model. **(A)** Total immune cells were FACS-sorted from the lungs of *MMTV-PyMT ^+/-^* at 9 weeks after the onset of primary breast cancer or control *MMTV-PyMT ^-/-^* wild-type littermates and subjected to the scRNA-seq analyses. **(B)** Neutrophils in the lungs of *MMTV-PyMT^+/-^* or control *MMTV-PyMT ^-/-^* wild-type littermates were examined by the FACS analyses. n = 10, mean ± SEM, * *p* < 0.05 (Student’s *t*-test). **(C to F)** Identification of immunosuppressive neutrophils in the metastatic lung tumors of *MMTV-PyMT ^+/-^* mouse model. Neutrophils were FACS-sorted from the peripheral blood, paracancerous lung tissues, and metastatic lung tumors of *MMTV-PyMT ^+/-^* mice at 9 weeks after the onset of primary breast cancer and subjected to the scRNA-seq analyses. **(C)** Neutrophil subtypes are defined in the pooled scRNA-seq data. Neu_Camp, Neu_Cxcr2, Neu_G0s2, and Neu_Entpd3, neutrophils with corresponding markers. **(D)** Differential distribution of neutrophil subtypes in the blood, paracancerous lung tissues, and metastatic lung tumors of *MMTV-PyMT ^+/-^*mice. **(E)** Pseudotime trajectory analysis of neutrophil subtypes defined in **(C)**. **(F)** Violin plot of the expression of immunosuppressive genes (*Cd274* and *Cd81*) and myeloid recruitment-related genes (*Il18*) in neutrophil subtypes. **(G)** Unique presence of immunosuppressive neutrophils in the metastatic lung tumors of *MMTV-PyMT ^+/-^* mouse model. Immunosuppressive neutrophils (CD45^+^ CD11b^+^ Ly6G^+^ Ly6C^-^ CD274^+^ ) in the indicated tissues of *MMTV-PyMT ^+/-^* at 9 weeks after the onset of primary breast cancer or control *MMTV-PyMT ^-/-^* littermates were examined by the FACS analyses. n = 5 for *MMTV-PyMT ^-/-^* mice and n = 3 for *MMTV-PyMT ^+/-^* mice. **(H)** High expression levels of CD274 in immunosuppressive neutrophils. *MMTV-PyMT* cancer cells and immunosuppressive neutrophils were FACS-sorted from the metastatic lung tumors of *MMTV-PyMT ^+/-^*mice and then subjected to the RNA-seq analyses together with *in vitro* cultured LLC cancer cells. Expression levels of *Cd274* in each cell type were determined. n = 4 for *MMTV-PyMT* cancer cells, n = 3 for LLC cancer cells, and n = 4 for immunosuppressive neutrophils, * *p* < 0.05 (one-way ANOVA test). **(I)** Enrichment of immunosuppressive neutrophils correlates with poor prognosis of human lung adenocarcinoma. The tumor samples with high enrichment scores (0.67 ∼ 0.90) *vs.* low enrichment scores (0.33 ∼ 0.50) in the TCGA dataset were used to compare the survival of patients (log-rank test).

Notably, Cd274/PD-L1 expressed by cancer cells is often considered the key blockage of antitumor immunity. We assessed the transcriptomic profiles of cancer cells and immunosuppressive neutrophils by the bulk RNA-seq analyses. It turned out to be a surprise that *Cd274* levels in immunosuppressive neutrophils were >100 times higher than those in LLC or *MMTV-PyMT* cancer cells (Fig. 3H), implicating immunosuppressive neutrophils as an indispensable component in designating the immune microenvironment of those lung tumors. To support the disease relevance of immunosuppressive neutrophils, we analyzed their signature genes in human lung adenocarcinoma samples in the TCGA database. It became evident that the enrichment of immunosuppressive neutrophils correlated with poor prognosis of lung adenocarcinoma patients (Fig. 3I). These results have identified the critical involvement of immunosuppressive neutrophils in lung tumors.

We explored the functional link between sympathetic signals and immunosuppressive neutrophils in lung tumors. The sympathetic neurotransmitter norepinephrine (NE) acts via the family of adrenergic receptors. The scRNA-seq data of neutrophil populations in the *MMTV-PyMT* cancer model revealed the exclusive expression of the β2-adrenergic receptor (Adrb2) among the family members (Fig. 4A). This distinct pattern of *Adrb2* expression in immunosuppressive neutrophils was confirmed by the qPCR analyses (Fig. 4B), raising the possibility that sympathetic signals might directly regulate the function of this neutrophil subtype. Indeed, the *in vitro* NE treatment of FACS-sorted immunosuppressive neutrophils effectively up-regulated the expression levels of their signature genes, such as *Cd274* and *Il18* (Fig. 4C). We further validated this sympathetic control of immunosuppressive neutrophils in the mouse cancer models. Cd274^+^ immunosuppressive neutrophils were evidently observed in the LLC lung tumors, but they became reduced in the condition of local sympathetic ablation by 6-OHDA (Fig. 4D and 4E). Similarly, the density of Cd274^+^ immunosuppressive neutrophils in metastatic lung tumors was significantly decreased in the 6-OHDA-treated *MMTV-PyMT ^+/-^* mice (Fig. 4F and 4G). Moreover, we tested the pharmacologic blockage of sympathetic signals in the context of immunotherapy. Lightsheet 3D imaging showed that while the administration of an anti-PD-L1 antibody alone inhibited tumor growth in the lungs of *MMTV-PyMT ^+/-^* mice, a detectable amount of tumor metastases still remained (Fig. 4H). In contrast, combining the anti-PD-L1 immunotherapy with an Adrb2 antagonist could effectively, or almost entirely in some cases, eliminate those lung metastases (Fig. 4H and 4I). These results have together elucidated an essential role of sympathetic modulation of the immunosuppressive neutrophils to promote lung tumors.

**Figure 4.**
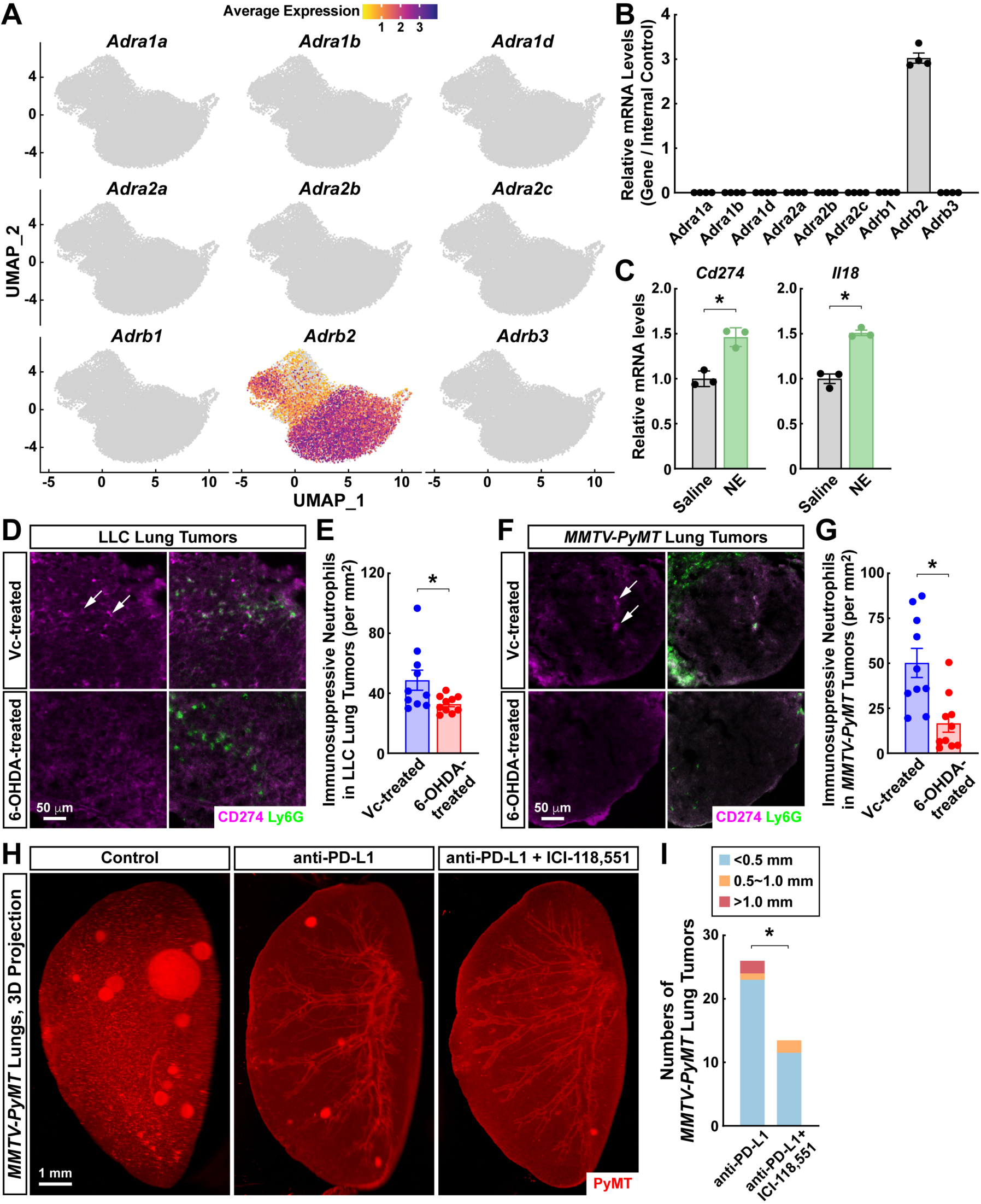
Sympathetic signals directly promote immunosuppressive neutrophils via Adrb2. **(A and B)** Distinct expression of *Adrb2* in neutrophils. **(A)** The average expression of adrenergic receptors was projected onto the UMAP plot of neutrophil subtypes defined in Fig. 3C. **(B)** Immunosuppressive neutrophils were FACS-sorted from the metastatic lung tumors of *MMTV-PyMT ^+/-^*mice 9 weeks after the onset of primary breast cancer. Expression levels of adrenergic receptors were determined by the qPCR analyses. n = 4. **(C)** The sympathetic neurotransmitter norepinephrine directly promotes the function of immunosuppressive neutrophils. Immunosuppressive neutrophils were FACS-sorted from the metastatic lung tumors of *MMTV-PyMT ^+/-^* mice and then *in vitro* treated with norepinephrine (NE). mRNA levels of *Cd274* and *Il18* were examined by the qPCR analyses. n = 3, mean ± SEM, * *p* < 0.05 (Student’s *t*-test). **(D to G)** Ablation of sympathetic signals abolishes immunosuppressive neutrophils in lung tumors. **(D and E)** The wild-type mice were intranasally treated with 6-OHDA or control vehicle [PBS containing 0.01% vitamin C (Vc)] and then inoculated with LLC cancer cells. **(D)** Representative images of the lung tumors processed by conventional tissue sectioning and CD274/Ly6G co-immunostaining were shown. **(E)** Numbers of immunosuppressive neutrophils in the lung tumors of Vc-treated or 6-OHDA-treated mice were quantified. n = 10, mean ± SEM, * *p* < 0.05 (Student’s *t*-test). **(F and G)** *MMTV-PyMT ^+/-^* female mice at 1 week after the onset of primary breast cancer were intranasally treated with 6-OHDA or control vehicle. **(F)** Representative images of the metastatic lung tumors processed by conventional tissue sectioning and CD274/Ly6G co-immunostaining were shown. **(G)** Numbers of immunosuppressive neutrophils in the metastatic lung tumors of Vc-treated or 6-OHDA-treated *MMTV-PyMT ^+/-^* mice were quantified. n = 10, mean ± SEM, * *p* < 0.05 (Student’s *t*-test). **(H and I)** Blockage of the Adrb2 signal potentiates the anti-PD-L1 immunotherapy. *MMTV-PyMT ^+/-^* female mice at 7 weeks after the onset of primary breast cancer were intravenously injected with the anti-PD-L1 antibody once per week without or with the daily treatment of the Adrb2 antagonist ICI-118,551. The intact lung tissues (left lobes) of the mice at 9 weeks after the onset of primary breast cancer were processed through the whole-tissue PyMT-immunolabeling. **(H)** Representative 3D-projection images at 1.26× magnification of the lightsheet imaging were shown. **(I)** Numbers and sizes of the metastatic lung tumors of *MMTV-PyMT ^+/-^* mice under indicated conditions were counted. 10 mice per condition, * *p* < 0.05 (two-way ANOVA test).

It has been well-documented that specific chemokines, including Cxcl1 and Cxcl2, regulate neutrophil chemotaxis in mice ^46,47^. Consistent with the neutrophil response in the lung tumors of LLC or *MMTV-PyMT* mouse models, we detected high levels of *Cxcl1* in both cancer cell types by the bulk RNA-seq results. In addition, *MMTV-PyMT* but not LLC cancer cells expressed *Cxcl2* among the CXCL class chemokines (Fig. 5A and 5B). Meanwhile, such transcriptomic data unexpectedly revealed the disparate capability of responding to sympathetic signals by these two cancer types (Fig. 5C). In particular, *MMTV-PyMT* cancer cells expressed multiple adrenergic receptors, including Adra2c, Adrb1, and Adrb2. On the other hand, LLC cancer cells had no detectable expression of the family members. Also, we found that the *in vitro* NE treatment of FACS-sorted MMTV-PyMT cancer cells increased *Cxcl1* and *Cxcl2* levels, and the Adrb2 agonist formoterol could recapitulate this sympathetic action (Fig. 5D). In contrast, the same treatment of LLC cancer cells with NE or formoterol did not affect those chemokines (Fig. 5E). In light of this finding, we examined the sympathetic ablation on neutrophil recruitment in the metastatic lung tumors of *MMTV-PyMT ^+/-^* mice. *Cxcl1* levels were markedly suppressed in the lung tumors of 6-OHDA-treated mice compared to the control condition, though *Cxcl2* expression exhibited high variations among tissue samples (Fig. 5F). Nevertheless, neutrophil accumulation in the *MMTV-PyMT* lung tumors could be mitigated by sympathetic ablation (Fig. 5G and 5H).

**Figure 5.**
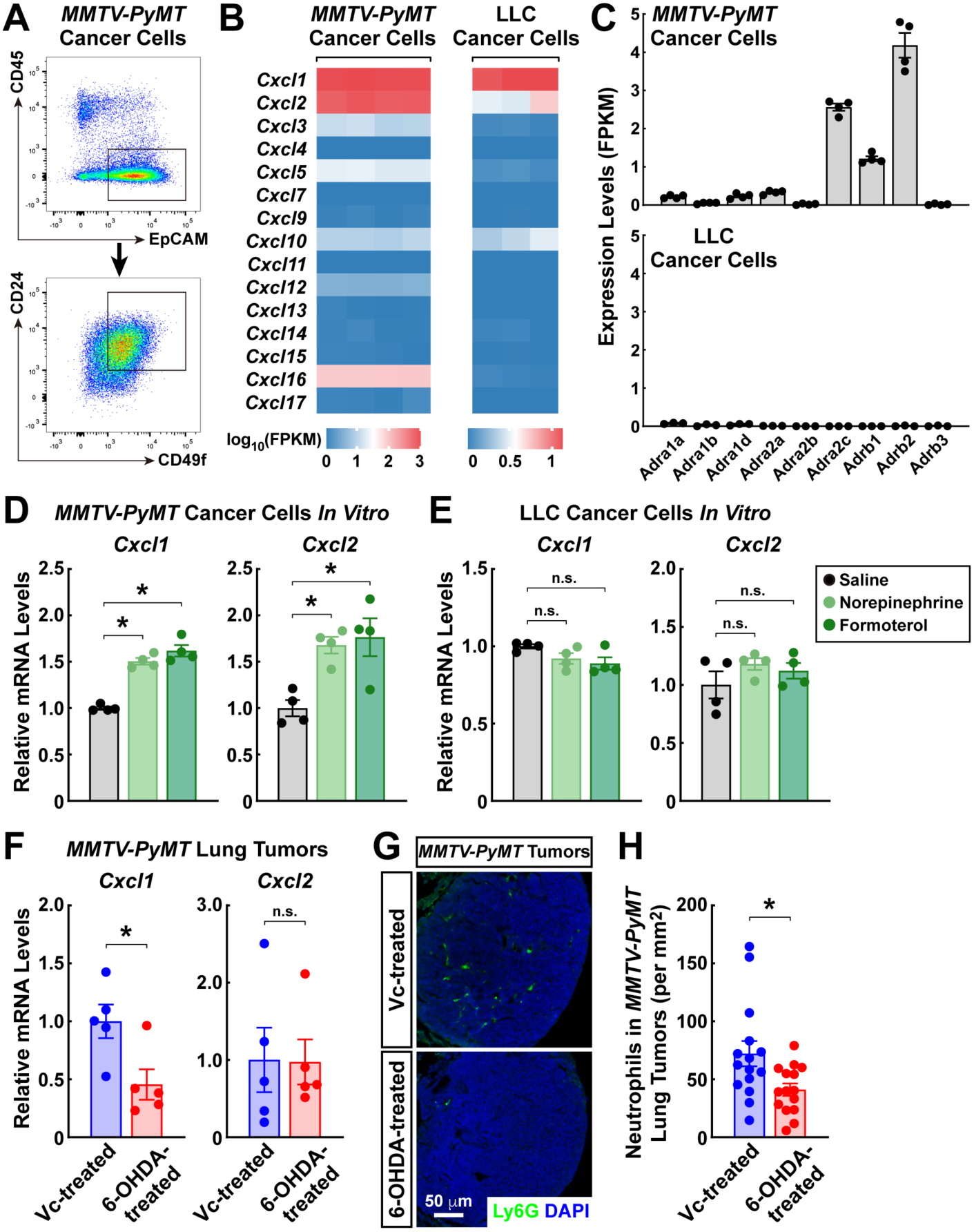
Sympathetic signals stimulate the specific type of cancer cells to recruit neutrophils. **(A to C)** Differential expression profiles of *MMTV-PyMT vs.* LLC cancer cells. **(A)** FACS diagram of primary cancer cells (EpCAM^+^ CD24^+^ CD49f^+^ CD45^-^ ) from the metastatic lung tumors of *MMTV-PyMT ^+/-^* mice. **(B and C)** Expression levels of the CXCL class chemokines **(B)** or adrenergic receptors **(C)** in *MMTV-PyMT vs.* LLC cancer cells were determined by the RNA-seq analyses. n = 4 for *MMTV-PyMT* cancer cells and n = 3 for LLC cancer cells. **(D and E)** Sympathetic signals stimulate the specific type of cancer cells to express Cxcl1 and Cxcl2. Primary *MMTV-PyMT* cancer cells **(D)** or LLC cancer cells **(E)** were *in vitro* treated with norepinephrine or the Adrb2 agonist formoterol. mRNA levels of *Cxcl1* and *Cxcl2* were examined by the qPCR analyses. n = 4, mean ± SEM, * *p* < 0.05, n.s., not significant (one-way ANOVA test). **(F to H)** Loss of local sympathetic signals blocks the neutrophil recruitment in lung tumors. *MMTV-PyMT^+/-^* female mice at 1 week after the onset of primary breast cancer were intranasally treated with 6-OHDA or control vehicle [PBS containing 0.01% vitamin C (Vc)]. **(F)** mRNA levels of *Cxcl1* and *Cxcl2* in the metastatic lung tumors of Vc-treated or 6-OHDA-treated *MMTV-PyMT ^+/-^* mice at 9 weeks after the onset of primary breast cancer were determined by the qPCR analyses. n = 5, mean ± SEM, * *p* < 0.05, n.s., not significant (Student’s *t*-test). **(G)** Representative images of the lung tumors processed by conventional tissue sectioning and Ly6G-immunostaining were shown. **(H)** Numbers of total neutrophils in the metastatic lung tumors of Vc-treated or 6-OHDA-treated *MMTV-PyMT ^+/-^* mice were quantified. n = 15, mean ± SEM, * *p* < 0.05 (Student’s *t*-test).

Notably, the loss of sympathetic innervations in the lung did not affect the neutrophil responses occurring in the spleen or peripheral blood of 6-OHDA-treated *MMTV-PyMT^+/-^* mice (Fig. S1C and S1D), supporting the specificity of sympathetic action to lung tumors. These results have illustrated the sympathetic stimulation of neutrophil recruitment as an additional layer of regulation on the immune microenvironment of specific lung tumors.

## Discussion

We reported for the first time the frequent engagement of sympathetic inputs in human NSCLC, as well as in the mouse models of lung tumors. This finding has suggested a direct connection of the tumor immune microenvironment to the body’s nervous system, substantiating clinical observations of the sympathetic influence on cancer prognosis ^33–35^. Meanwhile, whether human or mouse lung tumors receive other efferent neural inputs, particularly of parasympathetic origin, warrants future research. On the other hand, the existence of afferent sensory innervations within NSCLC is yet to be determined, which may act to relay immune cues derived from the tumor microenvironment to specific neurocircuits in the brain. Notably, the unparalleled power of whole-tissue immunolabeling and 3D imaging techniques is readily applicable to other cancer types, e.g., liver cancer, pancreatic cancer, or intestinal cancer, to precisely assess the neural structures in different human tumors. Indeed, our colleagues and we exploited 3D imaging to document neuroanatomy of the human liver ^48^, pancreas ^49^, or gastrointestinal tract ^50^, supporting the possibility of neural actions in tumorigenesis or tumor progression in those organs.

We identified a unique subtype of neutrophils exclusively present in the mouse lung tumors but absent in the peripheral blood or normal tissues. In contrast to their classic pro-inflammatory functions, this neutrophil subtype simultaneously possessed several immunosuppressive signals. This finding echoes the previous reports on the heterogenous neutrophil populations in human or mouse lung tumors, some of which correlated with the establishment of immunosuppressive microenvironment ^38,39,51^. More importantly, we elucidated that those immunosuppressive neutrophils specifically expressed the Adrb2 receptor, enabling their direct responsiveness to the sympathetic neurotransmitter norepinephrine. Whether this neutrophil subtype may be modulated by other neurotransmitters (e.g., glutamate or acetylcholine) or neuropeptides (e.g., calcitonin gene-related peptide or substance P) awaits further analyses. It appears conceivable that functional links between diverse neural signals and neutrophils can collectively designate the tumor immune microenvironment in the lung.

In addition to lung tumors, studies by colleagues and us have recently defined immunosuppressive neutrophil subtypes in liver cancer ^44^, prostate cancer ^52^, and glioma^45^. In light of the current study, an exciting question emerges as to whether the sympathetic regulation of immunosuppressive neutrophils may represent a common mechanism in other cancer types. Comprehensive profiling of adrenergic receptors in neutrophil subtypes of different human tumors by bulk or single-cell transcriptomics will provide an answer. Furthermore, we showed that the Adrb2 antagonist potentiated the therapeutic effect of anti-PD-L1 against mouse lung tumors, recapitulating the beneficial role of beta blockers reported in NSCLC patients ^35^. Whether targeting the Adrb2 signal in neutrophils can become a general adjuvant strategy for enhancing available immunotherapies warrants clinical investigations.

In sum, our study has demonstrated a critical aspect of sympathetic signals in designating the immune microenvironment of lung tumors, which has broad implications for understanding the complexity of cancer neuroscience.

## Acknowledgments

This work has been supported by the National Key Research and Development Program of China (2019YFA0802003 to J.Y.; #2022YFA1103900 to J.C.), the National Natural Science Foundation of China (#82173386 to M.Q.; #92059203 and #92259303 to K.C.; #32125017 and #32150008 to J.Y.), the Beijing Natural Science Foundation (#7234378 to Y.C.; #7232086 to J.Y.), the China Postdoctoral Science Foundation (#2022M710219 to Y.C.), the Chinese Academy of Medical Sciences (#2021RU002 and #2022-I2M-C&T-B-120 to K.C.; #2019-I2M-5-015 to J.C.), and Peking University People’s Hospital Scientific Research Development Funds (#RZ2022-04 to M.Q.; #RZ2022-03 to K.C.). Additional funds to K.C. have been from the Research Unit of Intelligence Diagnosis and Treatment in Early Non-Small Cell Lung Cancer. J.C. has been sponsored by the Changping Laboratory. J.Y. has been supported by the Center for Life Sciences at Peking University, the State Key Laboratory of Membrane Biology at Peking University, and Peking University Third Hospital Cancer Center. The authors declare no conflict of interest.

## Author Contributions

J.L., Y.C., J.C., and J.Y. performed experiments and analyzed data; M.Q., W.W., F.Y., and K.C. collected human lung tumor samples; K.C., J.C., and J.Y. designed the project and prepared the manuscript.

## Materials and Methods

Further information and requests for resources and reagents should be directed to and will be fulfilled by the Lead Contact, Jing Yang.

### Human lung cancer tissues

Human NSCLC samples were collected in compliance with the protocol approved by the Institutional Ethics Committees of Peking University People’s Hospital (#2020PHB009-01). Informed consent was signed by each involved patient.

### Mouse information and procedures

All the experimental procedures in mice were performed in compliance with the protocol approved by the Institutional Animal Care and Use Committees (IACUC) of Peking University and Chinese Institute for Brain Research. Mice were maintained on the 12hr/12hr light/dark cycle (light period 7:00 am ∼ 7:00 pm), with the chow diet and water available *ad libitum*. C57BL/6 wild-type female mice were purchased from Charles River International. *MMTV-PyMT ^+/-^* male mice on the FVB background were obtained from the Jackson Laboratory (#002374, RRID:IMSR_JAX:002374) and in-house bred with FVB wild-type female mice to generate *MMTV-PyMT ^+/-^* female littermates for the experiments.

To monitor the tumor onset, *MMTV-PyMT ^+/-^* female mice of 5 weeks were examined every other day for palpable mammary lumps. Tumor dimensions were measured every week, and tumor volumes were calculated as width (mm) × width (mm) × length (mm) / 2. Unless otherwise specified, the mice were euthanized for tissue harvesting and analyses at 9 weeks after the tumor onset.

For the establishment of LLC orthotopic lung tumors, C57BL/6J wild-type female mice of 8 weeks were intravenously injected with LLC cancer cells (1 × 10^6^ cells per mouse) in 200 μl of sterile phosphate-buffered saline (PBS). The mice were euthanized for tissue harvesting and analyses 30 days after the tumor inoculation.

For the measurement of plasma glucose levels, blood samples were collected at 9:00 am via tail bleeding and analyzed by Breeze2 glucometer (Bayer).

For the intranasal administration of 6-hydroxydopamine (6-OHDA; Sigma-Aldrich), *MMTV-PyMT ^+/-^*female mice at 1 week after the tumor onset were briefly anesthetized with 3% isoflurane. 500 μg of 6-OHDA dissolved in 50 μl of sterile PBS containing 0.01% vitamin C was intranasally delivered to each mouse every 48 hr for three times. For the control treatment, each mouse was intranasally administered 50 μl PBS containing 0.01% vitamin C.

For the anti-PD-L1 treatment, *MMTV-PyMT ^+/-^* female mice at 7 weeks after the tumor onset were intravenously injected with the anti-PD-L1 antibody (BioXcell, #BE0101, RRID:AB_10949073, diluted in sterile PBS at 1 mg/ml) at 5 mg/kg of body weight once per week.

For the administration of the Adrb2 antagonist, *MMTV-PyMT ^+/-^* female mice at 7 weeks after the tumor onset were daily treated with ICI-118,551 (MedChemExpress, diluted in sterile PBS at 1 mg/ml) at 5 mg/kg of body weight via intraperitoneal injection.

### Whole-tissue immunolabeling and optical clearing

The whole-tissue immunolabeling of unsectioned human or mouse tissues was performed according to the iDISCO(+) technique ^53,54^ with several modifications. Surgically resected human lung cancer tissues were immediately fixed in PBS / 1% paraformaldehyde (PFA) / 10% sucrose at room temperature for 1 hr. The tissues were further fixed in PBS / 1% PFA at 4°C overnight and washed with PBS at room temperature for 1 hr three times. Alternatively, *MMTV-PyMT ^+/-^*female mice of indicated conditions were anesthetized and perfused sequentially with 20 ml PBS / 100 μg/ml heparin and 20 ml PBS / 1% PFA / 10% sucrose / 100 μg/ml heparin. Mouse lung tissues were dissected out, post-fixed in PBS / 1% PFA at room temperature for 2 hr, and washed with PBS at room temperature for 1 hr three times. All the incubation steps were performed with gentle rotation.

The PFA-fixed human or mouse tissues were incubated at room temperature with 20% methanol (diluted in ddH_2_O) for 1 hr, 40% methanol for 1 hr, 60% methanol for 1 hr, 80% methanol for 1 hr, and 100% methanol for 1 hr twice. The tissues were decolorized with a mixture of 30% H_2_O_2_ and 100% methanol (v:v = 1:10) at 4°C for 48 hr. The tissues were then incubated at room temperature with 80% methanol for 1 hr, 60% methanol for 1 hr, 40% methanol for 1 hr, 20% methanol for 1 hr, and PBS / 0.2% TritonX-100 / 10 mM EDTA-Na (pH 8.0) / 10% DMSO for 1 hr. The tissues were further permeabilized in PBS / 0.2% TritonX-100 / 0.1% deoxycholate / 10 mM EDTA-Na (pH 8.0) / 10% DMSO at room temperature overnight. All the incubation steps were performed with gentle rotation.

The human or mouse tissues were blocked with PBS / 0.2% Tween-20 / 10 μg/ml heparin / 5% normal donkey serum / 5% DMSO at room temperature overnight. The tissues were immunolabeled with the primary antibodies diluted (final concentration of 2 μg/ml) in PBS / 0.2% Tween-20 / 10 μg/ml heparin / 5% normal donkey serum / 5% DMSO at 37°C for 72 hr. The primary antibodies used in this study were rabbit anti-tyrosine hydroxylase (TH; Millipore, #AB152, RRID:AB_390204) and rat anti-polyomavirus middle T antigen (PyMT; Santa Cruz, #sc53481, RRID:AB_630138). The tissues were washed with PBS / 0.2% Tween-20 / 10 μg/ml heparin at 37°C for 12 hr, with the fresh buffer changed every 2 hr. The tissues were further immunolabeled with the secondary antibodies diluted (final concentration of 4 μg/ml) in PBS / 0.2% Tween-20 / 10 μg/ml heparin / 5% normal donkey serum / 5% DMSO at 37°C for 72 hr. The secondary antibodies used in this study were Alexa Fluor 647-conjugated donkey anti-rabbit IgG (Thermo Fisher Scientific, #A31573, RRID:AB_2536183) and Alexa Fluor 568-conjugated donkey anti-rat IgG (Thermo Fisher Scientific, #A78946, RRID:AB_2910653). The tissues were washed with PBS / 0.2% Tween-20 / 10 μg/ml heparin at 37°C for 24 hr, with the fresh buffer changed every 4 hr. All the incubation steps were performed with gentle rotation.

The immunolabeled human or mouse tissues were washed in PBS at room temperature for 2 hr and embedded in PBS / 0.8% agarose for the optical-clearing steps. The tissue blocks were incubated at room temperature with 20% methanol (diluted in ddH_2_O) for 1 hr three times, 40% methanol for 1 hr, 60% methanol for 1 hr, 80% methanol for 1 hr, 100% methanol for 1 hr, and 100% methanol overnight. The tissue blocks were then incubated at room temperature with a mixture of dichloromethane and methanol (v:v = 2:1) for 3 hr, followed by 100% dichloromethane for 1 hr three times. The tissue blocks were finally incubated at room temperature with 100% dibenzyl-ether for 12 hr twice. All the incubation steps were performed with gentle rotation.

### Lightsheet microscopy and 3D reconstruction

The optically-cleared human or mouse tissues were imaged on the LaVision Biotec Ultramicroscope II equipped with the 2×/NA0.5 objective covered with a 10mm-working-distance dipping cap. The tissues were immersed in the imaging chamber filled with 100% dibenzyl-ether. For imaging at 1.26× magnification (0.63× zoom) or 2.0× magnification (1.0× zoom), each tissue was scanned by three combined lightsheets from the left side with a step size of 5 μm. For imaging at 8.0× magnification (4.0× zoom), each tissue was scanned by a single lightsheet (middle position) from the left side with a step size of 1 μm.

Imaris (https://imaris.oxinst.com/packages) was used to 3D reconstruct the image stacks obtained from lightsheet microscopy. TH-positive sympathetic innervations in human lung cancer tissues or the diameters of PyMT-positive tumors in mouse lung tissues were manually assessed. Orthogonal projections were generated from the 3D reconstructed stacks for representative images shown in the figures. For the display purpose, a gamma correction of 0.9 ∼ 1.2 was applied to the raw data.

### Fluorescence-activated cell sorting (FACS)

The spleens were freshly dissected from the mice of indicated conditions and cut into small pieces on ice. The tissues were mashed through a 70-μm cell strainer in PBS / 2% heat-inactivated fetal bovine serum (HI-FBS; Sigma-Aldrich) / 10 mM EDTA-Na (pH 8.0), and cell suspensions were centrifuged at 500 *g* for 5 min. The cells were re-suspended in the ammonium-chloride-potassium (ACK) buffer (Thermo Fisher Scientific) to lyse red blood cells and centrifuged at 500 *g* for 5 min. The cells were then re-suspended in PBS / 2% HI-FBS / 10 mM EDTA-Na (pH 8.0) and stained with the intended FACS antibodies.

The blood samples were collected from the mice of indicated conditions via tail bleeding into PBS / 2% HI-FBS / 10 mM EDTA-Na (pH 8.0). The samples were centrifuged at 500 *g* for 5 min and re-suspended in the ACK buffer to lyse red blood cells. The cells were then centrifuged at 500 *g* for 5 min, re-suspended in PBS / 2% HI-FBS / 10 mM EDTA-Na (pH 8.0), and stained with the intended FACS antibodies.

The lung tissues were freshly dissected from the mice of indicated conditions and cut into small pieces on ice. The tissues were digested in RPMI 1640 (Thermo Fisher Scientific) / 0.1 mg/ml Liberase TL (Roche) / 20 μg/ml DNase I (Sigma-Aldrich) / 10 mM HEPES / 3% HI-FBS at 37°C for 15 min. The tissues were mashed through a 70-μm cell strainer and centrifuged at 500 *g* for 5 min. The cells were re-suspended in the ACK buffer to lyse red blood cells. The cells were centrifuged again at 500 *g* for 5 min, re-suspended in PBS / 2% HI-FBS / 10 mM EDTA-Na (pH 8.0), and stained with the intended FACS antibodies.

FACS-stained immune cells were analyzed on the BD LSRFortessa, and the results were processed by FlowJo (https://www.flowjo.com). Alternatively, FACS-stained tumor cells or neutrophils were sorted on the Beckman MoFlo Astrios EQ.

### Single-cell RNA sequencing (scRNA-seq)

Total immune cells (CD45^+^) or neutrophils (CD45^+^ CD11b^+^ Ly6G^+^ Ly6C^-^ ) were FACS-sorted from the blood, lung tissues, or tumors of *MMTV-PyMT ^+/-^* female mice and subjected to scRNA-seq by the 10x Genomics or the Singleron. The scRNA-seq data were deposited to the Sequence Read Archive (https://www.ncbi.nlm.nih.gov/sra): SRR25435720 for total immune cells in the lung tissues of wild-type FVB female mice; SRR25445370 for total immune cells in the lung tissues of *MMTV-PyMT ^+/-^* female mice; SRR25472801 for neutrophils in the blood of *MMTV-PyMT ^+/-^* female mice; SRR25488388 for neutrophils in the tumors of *MMTV-PyMT ^+/-^*female mice.

10x Genomics data preprocessing was performed according to the manufacturer’s instructions (https://support.10xgenomics.com/single-cell-gene-expression/software/pipelines/latest/advanced/references), and CellRanger (v7.0.1) was used for sequence alignment and gene expression quantification. Singleron data preprocessing was performed by CeleScope (v1.13.0) (https://github.com/singleron-RD/CeleScope). Both types of data preprocessing used mm10 as the reference mouse genome. We filtered out the genes expressed in <10 cells, as well as the cells expressing <200 genes, >10% mitochondrial reads, or >20% ribosomal genes. Solo in scvi-tools ^55^ was employed to remove doublets using the top 2,000 variant genes and default parameters.

We chose single-cell Variational Inference (scVI) ^56^ to integrate different scRNA-seq samples according to the software instructions (https://docs.scvi-tools.org/en/stable/tutorials/index.html). The integrated data were clustered by SCANPY ^57^, and the resulting H5ad files were converted into Seurat objects by Sceasy (https://github.com/cellgeni/sceasy) and run through Seurat v.4 ^58^ to visualize the gene markers for each cluster. We combined ScType ^59^ and PanglaoDB ^60^ for the initial determination of cell types. For the classification of neutrophil subtypes, we referred to the recent works by our colleagues and us ^45,61^.

TCGA databases of human lung adenocarcinoma were obtained from GDAC Firehose of Broad Institute (https://gdac.broadinstitute.org). GSVA ^62^ was used to calculate the enrichment score of immunosuppressive neutrophils in each sample. The tumor samples with high enrichment scores (0.67 ∼ 0.90) and the tumor samples with low enrichment scores (0.33 ∼ 0.50) in the dataset were used to compare the survival of patients.

### Bulk RNA sequencing (RNA-seq)

LLC cancer cells were cultured in Dulbecco’s Modified Eagle Medium (DMEM; Thermo Fisher Scientific) supplemented with 10% HI-FBS, 1× MEM non-essential amino acids (Gibco), 2 mM *L*-glutamine, 100 U/ml penicillin, and 100 μg/ml streptomycin. *MMTV-PyMT* cancer cells (EpCAM^+^ CD24^+^ CD49f^+^ CD45^-^ ) or immunosuppressive neutrophils (CD45^+^ CD11b^+^ Ly6G^+^ Ly6C^-^ CD274^+^ ) were FACS-sorted from the tumors of *MMTV-PyMT ^+/-^* female mice. Total RNAs of the cells were extracted by the RNeasy Mini Kit (Qiagen) and subjected to single-end RNA-seq by the Beijing Genomics Institute. Gene expression levels were normalized as fragments per kilobase per million (FPKM). The RNA-seq data were deposited to the Sequence Read Archive: SRR25535931, SRR25535932, and SRR25535933 for LLC cancer cells; SRR25244854, SRR25244855, SRR25244856 and SRR25244857 for *MMTV-PyMT* cancer cells; SRR25245226, SRR25245227, SRR25245228, and SRR25245229 for neutrophils.

### *In vitro* treatments

*MMTV-PyMT* cancer cells (EpCAM^+^ CD24^+^ CD49f^+^ CD45^-^ ) or immunosuppressive neutrophils (CD45^+^ CD11b^+^ Ly6G^+^ Ly6C^-^ CD274^+^ ) were FACS-sorted from the tumors of *MMTV-PyMT ^+/-^* female mice. *MMTV-PyMT* cancer cells were cultured in DMEM supplemented with 1% insulin-transferrin-selenium (Thermo Fisher Scientific), 100 U/ml penicillin, and 100 μg/ml streptomycin. Immunosuppressive neutrophils were cultured in RPMI 1640 supplemented with 10% HI-FBS, 1× MEM non-essential amino acids, 2 mM *L*-glutamine, 100 U/ml penicillin, and 100 μg/ml streptomycin. After resting at 37°C for 4 hr, *MMTV-PyMT* cancer cells or neutrophils were treated with a final concentration of 20 μM norepinephrine or 20 μM formoterol (Selleck) for 2 hr. Also, cultures of LLC cancer cells were subjected to the same treatment of norepinephrine or formoterol. Total RNAs of the cells were then extracted by the RNeasy Mini Kit and analyzed by the SYBR Green Real-Time PCR Kit (Thermo Fisher Scientific). *Cyclophilin* mRNA levels were utilized as the internal control.

### Immunofluorescence staining

*MMTV-PyMT ^+/-^*female mice of indicated conditions were anesthetized and perfused sequentially with 20 ml PBS / 100 μg/ml heparin and 20 ml PBS / 1% PFA / 100 μg/ml heparin. Mouse lung tissues were dissected out and post-fixed in PBS / 1% PFA at 4°C overnight. The tissues were then cryopreserved in PBS / 30% sucrose at 4°C overnight before 10-μm sectioning. The tissue sections were immunostained with rabbit anti-tyrosine hydroxylase (Millipore, #AB152, RRID:AB_390204), rat anti-Ly6G (eBioscience, #16-9668-82, RRID:AB_2573128), rat anti-CD8a (BioLegend, #100702, RRID:AB_312741), or rabbit anti-PD-L1 (Cell Signaling Technology, #64988, RRID:AB_2799672), followed by Alexa Fluor 488-conjugated donkey anti-rat IgG (Thermo Fisher Scientific, #A21208, RRID:AB_2535794) or Alexa Fluor 568-conjugated donkey anti-rabbit IgG (Thermo Fisher Scientific, #A10042, RRID:AB_2534017). The immunostained tissue sections were scanned by Axio Scan Z1, and the immune cell numbers within tumor regions were manually quantified in ImageJ (https://imagej.net/ij).

### Giemsa staining

Neutrophils were FACS-sorted from the blood, spleens, lung tissues, and lung tumors of LLC or *MMTV-PyMT ^+/-^* cancer models. Neutrophils were then spread on glass slides and stained with the Giemsa Stain Kit (Solarbio, #G4640). The stained samples were scanned under a bright field by Zeiss Axio Scan Z1 equipped with a 20× objective. Two hundred neutrophils were randomly selected from the images of each condition, and the number of nuclear lobes in each cell was manually counted.

### Statistical methods

Student’s *t*-tests (two-tailed unpaired) and ANOVA with *post hoc* tests were performed using GraphPad Prism 8.4.3 (http://www.graphpad.com/scientific-software/prism). Statistical details of the experiments are included in the figure legends.

---

**Figure S1.**
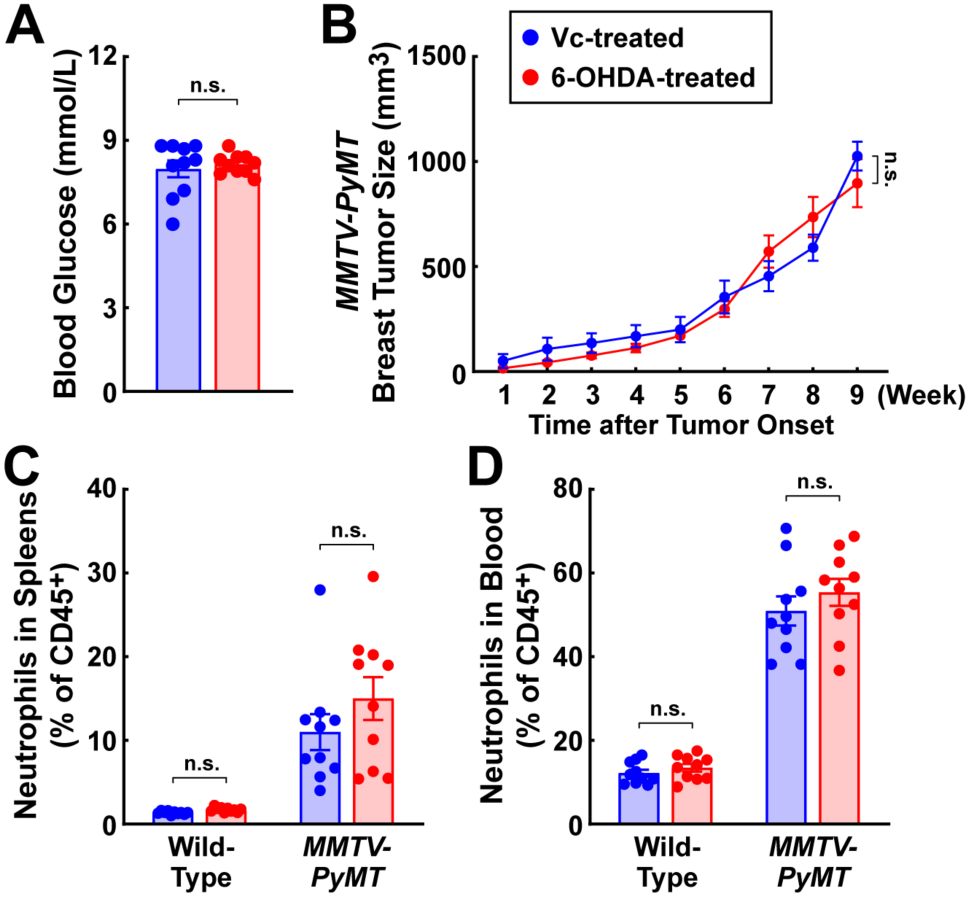
Ablation of local sympathetic inputs in the lung. Related to Figure 2. **(A)** The wild-type mice were subjected to the intranasal treatment of 6-OHDA or control vehicle [PBS containing 0.01% vitamin C (Vc)]. Blood glucose levels of the Vc-treated or 6-OHDA-treated mice were compared at 2 weeks after the intranasal instillation. n = 10, mean ± SEM, n.s., not significant (Student’s *t*-test). **(B to D)** *MMTV-PyMT ^+/-^*female mice at 1 week after the onset of primary breast cancer were intranasally treated with 6-OHDA or control vehicle [PBS containing 0.01% vitamin C (Vc)]. *MMTV-PyMT ^+/+^*female littermates were also subjected to the 6-OHDA or Vc treatment. **(B)** The volumes of primary breast tumors of the Vc-treated or 6-OHDA-treated *MMTV-PyMT ^+/-^* mice were monitored for 8 weeks after the intranasal instillation. n = 10, mean ± SEM, n.s., not significant (two-way ANOVA test). **(C and D)** Neutrophils in the spleens **(C)** or the blood **(D)** of Vc-treated or 6-OHDA-treated *MMTV-PyMT ^+/-^* and control *MMTV-PyMT ^-/-^* wild-type littermates were examined by the FACS analyses. n = 10, mean ± SEM, n.s., not significant (two-way ANOVA test).

**Figure S2.**
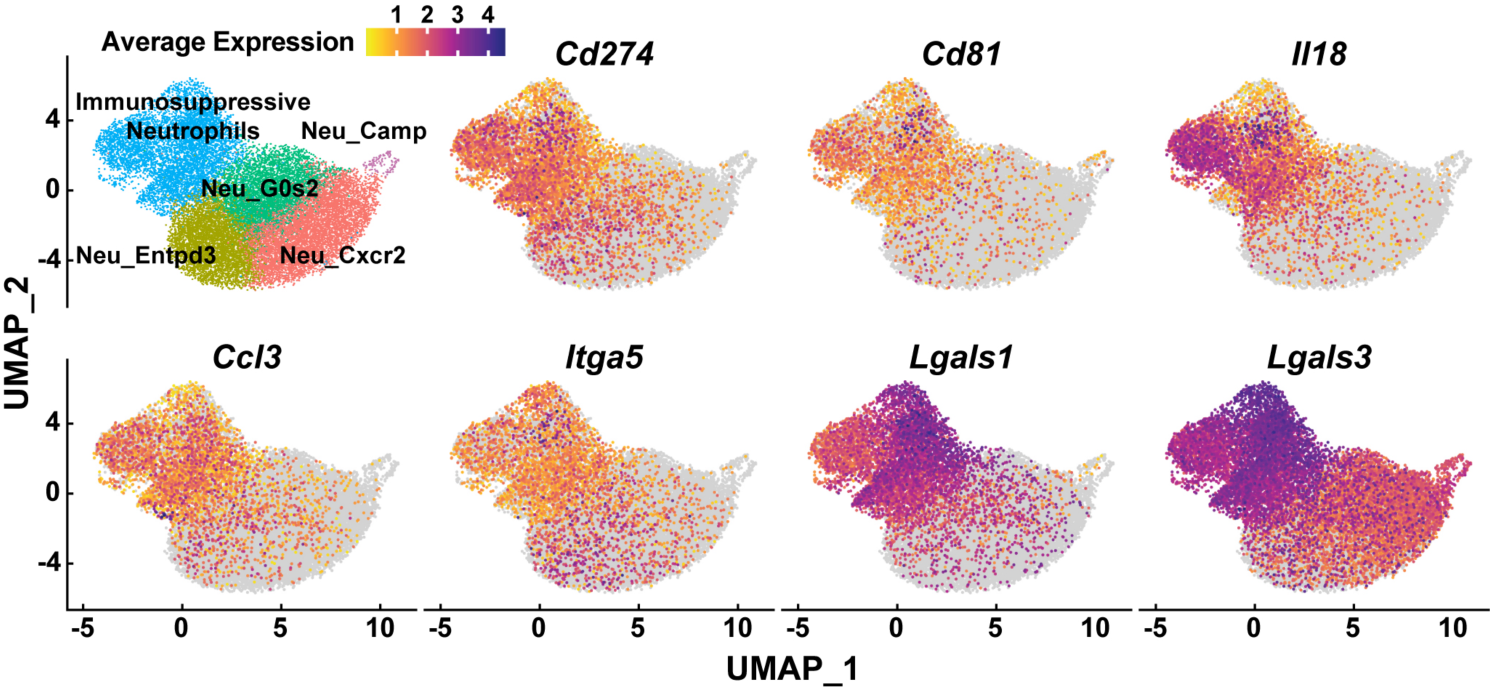
Identification of immunosuppressive neutrophils in the metastatic lung tumors of *MMTV-PyMT ^+/-^* mouse model. Related to Figure 3. Neutrophil subtypes are defined in the pooled scRNA-seq data of the peripheral blood, paracancerous lung tissues, and metastatic lung tumors of *MMTV-PyMT ^+/-^* female mice at 9 weeks after the onset of primary breast cancer. Neu_Camp, Neu_Cxcr2, Neu_G0s2, and Neu_Entpd3, neutrophils with corresponding markers. The average expression of immunosuppressive genes (*Cd274*, *Cd81*, *Itga5*, *Lgals1*, and *Lgals3*) and myeloid recruitment-related genes (*Ccl3* and *Il18*) was projected onto the UMAP plot of neutrophil subtypes.

**Figure S3.**
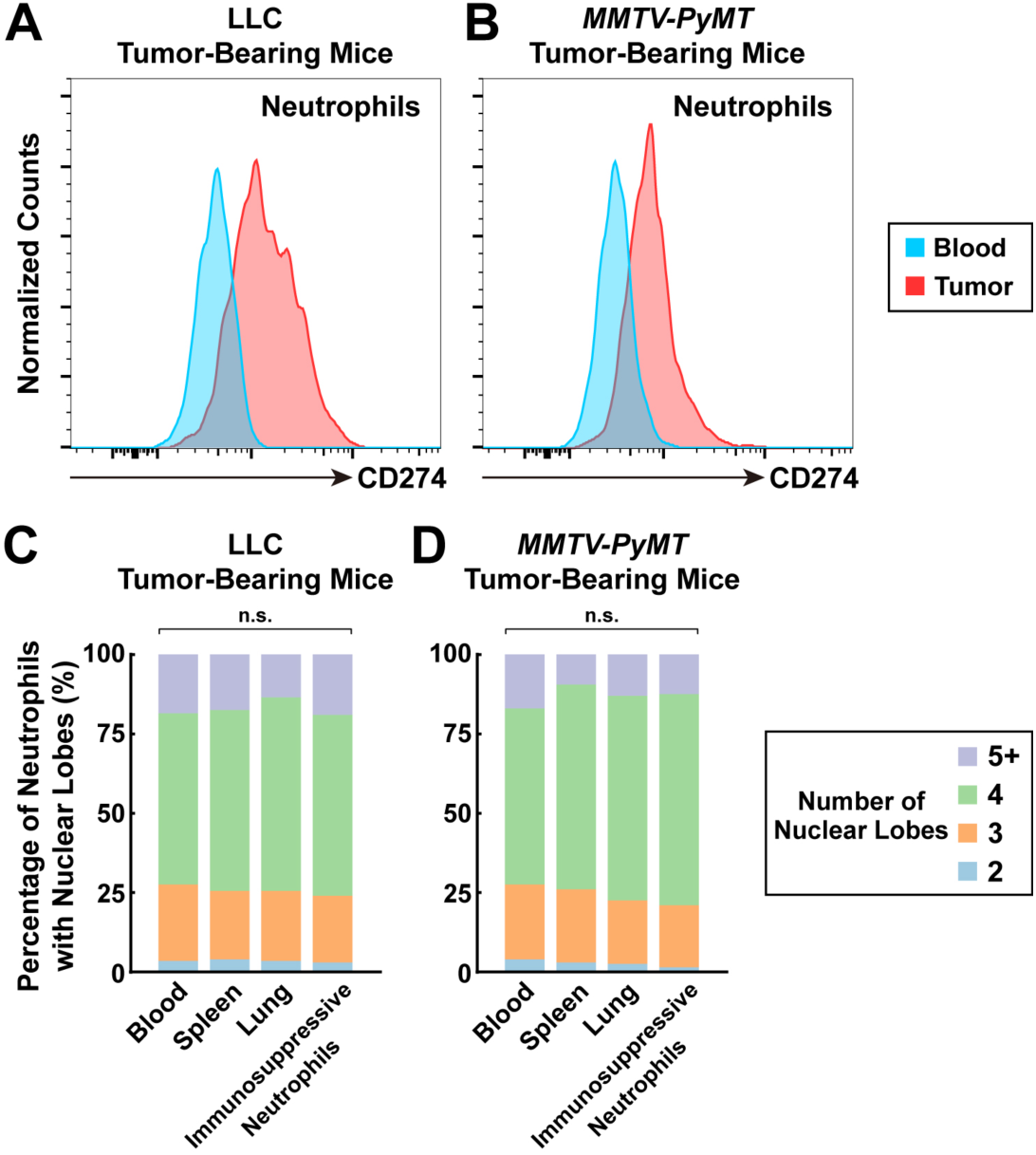
Characterization of immunosuppressive neutrophils in the mouse lung tumors. Related to Figure 3. **(A and B)** Expression levels of CD274 in the neutrophils of the blood and lung tumors of LLC tumor-bearing mice **(A)** or *MMTV-PyMT ^+/-^* mice **(B)** were assessed by the FACS analyses. **(C and D)** Nuclear segmentation of the neutrophils FACS-sorted from the blood, spleens, paracancerous lung tissues, and lung tumors of tumor-bearing LLC mice **(C)** or *MMTV-PyMT ^+/-^* mice **(D)** were examined by the Giemsa staining. 200 cells per condition, n.s., not significant (two-way ANOVA test).

**Table S1.** Information of NSCLC patients. Related to Figure 1.

| Patient Number | Diagnosis / TNM Classification | Gender | Age | Smoking History | Sympathetic Innervations in Tumor |
| --- | --- | --- | --- | --- | --- |
| 1 | Invasive adenocarcinoma, T2aN0M0, IB | F | 67 | No | ++++ |
| 2 | Invasive adenocarcinoma, T1bN0M0, IA2 | M | 68 | No | ++ |
| 3 | Invasive adenocarcinoma, T1cN0M0, IA3 | F | 63 | No | ++ |
| 4 | Invasive adenocarcinoma, T2aN0M0, IB | M | 52 | Yes | +++ |
| 5 | Invasive adenocarcinoma, T1aN0M0, IA1 | F | 65 | No | ++++ |
| 6 | Large-cell neuroendocrine carcinoma, T2aN0M0, IB | M | 70 | Yes | ++ |
| 7 | Invasive adenocarcinoma, T2N0M0, IB | F | 85 | No | ++ |
| 8 | Invasive adenocarcinoma, T2aN0M0, IB | M | 83 | Yes | n.d. |
| 9 | Squamous cell carcinoma, TbN0M0, IA2 | M | 68 | No | + |
| 10 | Invasive adenocarcinoma, T2bN0M0, IIA | M | 28 | No | + |
| 11 | Invasive adenocarcinoma, T1cN0M0, IA3 | M | 58 | Yes | n.d. |
| 12 | Invasive adenocarcinoma, T1N0M0, IA1 | M | 70 | No | + |
| 13 | Invasive adenocarcinoma, T1bN2M0, IIIA | M | 74 | No | + |
| 14 | Invasive adenocarcinoma, T2N0M0, IB | F | 46 | No | + |
| 15 | Invasive adenocarcinoma, T1cN0M0, IA3 | M | 60 | No | + |
| 16 | Squamous cell carcinoma, T2aN2M0, IIIA | M | 74 | Yes | ++ |
| 17 | Squamous cell carcinoma, T2bN0M1, IV | F | 66 | No | + |
| 18 | Invasive adenocarcinoma, T2aN0M0, IB | M | 69 | No | + |
| 19 | Invasive adenocarcinoma, T1aN0M0, IA1 | M | 71 | Yes | + |
| 20 | Invasive adenocarcinoma, T2aN0M0, IB | F | 63 | No | +++ |

++++, >50 sympathetic axons; +++, 25~50 axons; ++, 10~25 axons; +, <10 axons;
n.d., not detected.

